# AI-QuIC: Machine Learning for Automated Detection of Misfolded Proteins in Seed Amplification Assays

**DOI:** 10.1101/2024.10.16.618742

**Authors:** Kyle D. Howey, Manci Li, Peter R. Christenson, Peter A. Larsen, Sang-Hyun Oh

## Abstract

Advancements in AI, particularly deep learning, have revolutionized protein folding modeling, offering insights into biological processes and accelerating drug discovery for protein misfolding diseases. However, detecting misfolded proteins associated with neurodegenerative disorders, such as Alzheimer’s, Parkinson’s, ALS, and prion diseases, relies on Seed Amplification Assays (SAAs) analyzed through manual, time-consuming, and potentially inconsistent methods. We introduce AI-QuIC, an AI-driven platform that automates the analysis of Real-Time Quaking-Induced Conversion (RT-QuIC) assay data, a type of SAA crucial for detecting misfolded proteins. Utilizing a well-labeled RT-QuIC dataset of over 8,000 wells—the largest curated dataset for chronic wasting disease prion detection—we applied various AI models to classify true positive, false positive, and negative reactions. Notably, our deep-learning-based model achieved over 98% sensitivity and 97% specificity. By learning directly from raw fluorescence data, deep learning simplifies the SAA-analysis workflow. Automating and standardizing SAA data interpretation with AI-QuIC provides robust, scalable, and consistent diagnostic solutions.

## Introduction

Advancements in AI, in particular deep learning^1^, have revolutionized numerous scientific fields, including life sciences and disease diagnostics. Pioneering developments in protein structure prediction algorithms, such as AlphaFold^2^ and RosettaFold^3,4^ have demonstrated the impact of AI on predicting protein folding, with significant potential implications for accelerated drug discovery. These breakthroughs have set the stage for applying AI to tackle challenges in neurodegenerative diseases characterized by protein misfolding.

Early detection of misfolded proteins is crucial for diagnosing and managing neurodegenerative diseases such as Alzheimer’s, Parkinson’s, and amyotrophic lateral sclerosis (ALS). Seed amplification assays (SAAs), such as protein misfolding cyclic amplification (PMCA)^5^, quaking-induced conversion (QuIC)^6–8^, and real-time QuIC (RT-QuIC)^9^, have emerged as powerful tools for detecting these pathogenic proteins in various neurodegenerative diseases^10–16^. SAAs cyclically amplify protein misfolding to enable the rapid conversion of a large excess of the monomeric substrate into prion-like amyloid fibrils with minimal quantities of seeds formed by misfolded proteins^8,17^. These assays recapitulate the seeding-nucleation mechanisms of protein misfolding and help determine whether samples contain detectable (positive) or undetectable (negative) levels of misfolded proteins^5^. Misfolded proteins detected by SAAs include prions in prion diseases (e.g., Creutzfeldt-Jakob Disease)^6,9,14,18–20^, amyloid-beta^21^ and tau^10,11,22^ in Alzheimer’s disease, alpha-synuclein in synucleinopathies^12,15,23,24^, and TAR DNA-binding protein 43 in ALS^25^. By achieving ultrasensitive detection of misfolded proteins, SAAs hold great potential for early diagnosis and prognosis of such disorders^14^.

While SAAs have broad applications across multiple neurodegenerative diseases in humans, their development has benefited significantly from prion diseases in animals, especially Chronic Wasting Disease (CWD) that naturally occurs in cervids (e.g., deer, elk, and moose)^26–33^. Therefore, CWD is considered a robust model for studying protein misfolding in RT-QuIC.

Sharing the fundamental principles of SAAs, RT-QuIC assays are performed in multi-well plates with 4-8 technical replicates/reactions per sample^9^. The plate undergoes cyclic shaking and incubation while heated to facilitate fibril fragmentation and promote seeding-nucleation, resulting in the exponential amplification of misfolded proteins^9^. ThT, a rotor dye, exhibits enhanced fluorescence upon binding to growing amyloid fibrils in the reaction mixture due to restricted rotational motion, enabling real-time monitoring of amyloid formation^9,34^. Several methods have been applied to RT-QuIC data to establish predictable values using various metrics^29,35,36^ (Figure 1a), but it is unclear whether there is unutilized information in the raw fluorescence measurements. The complexity of interpreting amplification signals is compounded by factors such as the concentration of misfolded proteins, strain conformation, and cross-seeding^35,37,38^. In short, processes for the optimization of data interpretation and standardization across different assay platforms and disease-associated proteins remain unexplored.

**Fig. 1.**
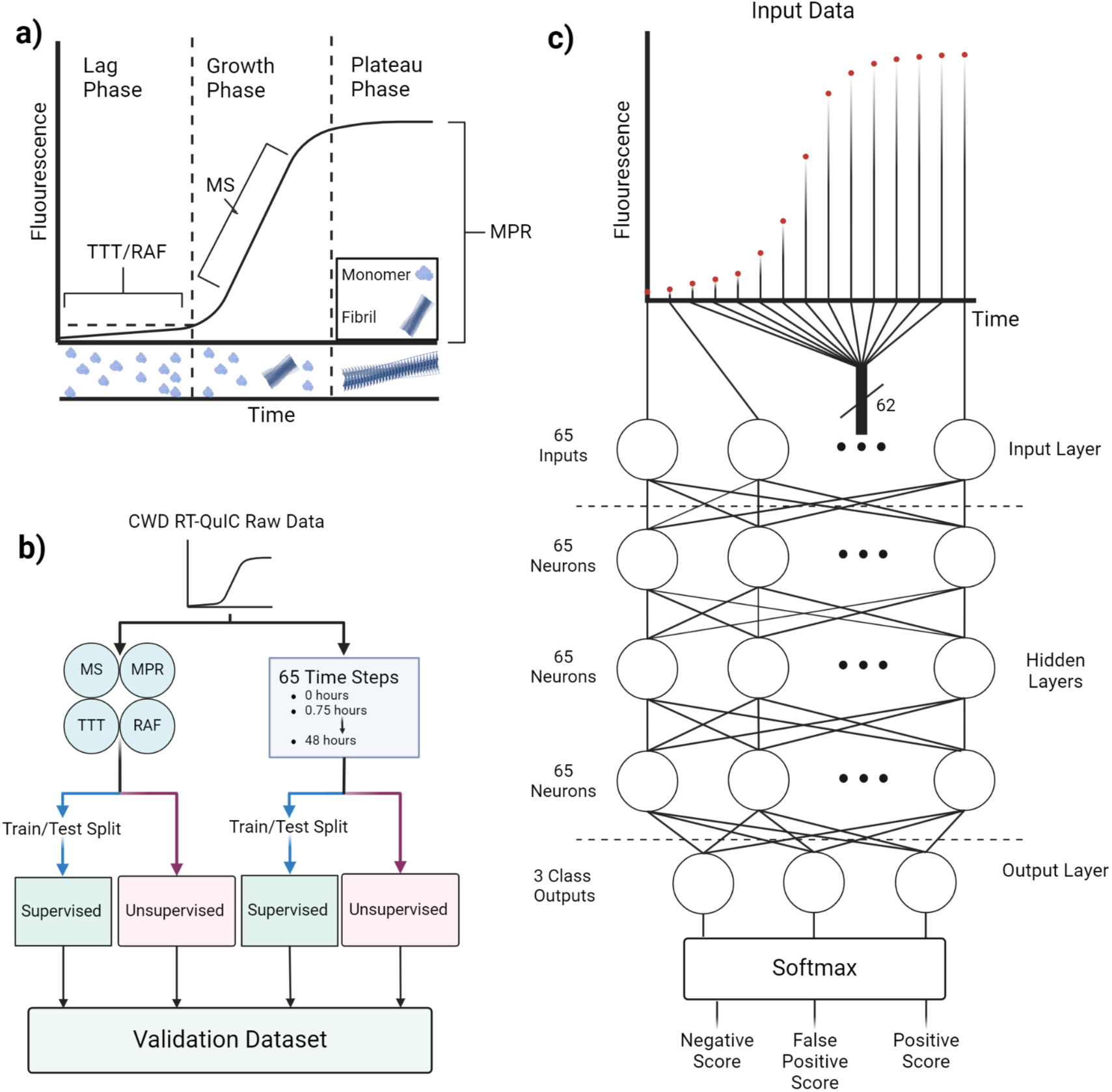
Visualization of Key Concepts. **a)** Example graph of the RT-QuIC data from a single reaction with the source of Time to Threshold (TTT), Rate of Amyloid Formation (RAF), Max Slope (MS), and Max Point Ratio (MPR) highlighted. The different phases of the reaction are identified with a visual of how the monomers form a fibril, increasing the fluorescence. **b)** Flowchart of how the data is processed in the study. **c)** Diagram of the multilayer perceptron used in this study with 65 input features. The network also includes 3 hidden layers, each with 65 neurons and a single output layer with 3 classes.

Drawing conclusions (positive vs. negative for seeding activity) from RT-QuIC reactions requires expertise by trained personnel, often requiring nuanced and subjective knowledge that is hard to formalize and articulate in human language or as a written manual^39^. AI is uniquely suited for such challenges as it can replicate and capture expert knowledge, identify complex patterns, and make accurate predictions, thereby enhancing the reliability and efficiency of RT- QuIC assays. Integrating AI into these processes bridges gaps in human expertise and enables robust, scalable testing operations.

The fluorescence measurements at each time point in an RT-QuIC reaction form an RT- QuIC curve, which can be used directly or summarized into derived metrics to serve as features or input parameters for training ML models. Many AI models exist, each with a unique approach to carrying out such tasks. For example, clustering classifiers such as K-Means are a subset of unsupervised AI models that identify patterns in data and assign labels associated with those patterns^40,41^. These can be used while considering whether an RT-QuIC reaction has seeding activity (positive) or not (negative), making them invaluable for use with large, unlabeled RT-QuIC datasets, as verifying labels can be time-consuming. More complex and supervised models, namely Support Vector Machines (SVMs) and Multilayer Perceptrons (MLPs), can be trained to identify positive RT-QuIC reactions on labeled data, potentially with better performance^42,43^. Investigating AI-driven automation and standardization techniques for interpreting SAA data, particularly from RT-QuIC assays, holds great promise in further enhancing the already significant impact of SAAs on the detection and diagnosis of neurodegenerative diseases.

The objective of this proof-of-concept study is to demonstrate the potential of AI in detecting the seeding activity of misfolded proteins and to explore the effectiveness of various AI models including deep learning models, such as MLPs (Figure 1c), in analyzing data from SAAs, using RT-QuIC data from CWD prions as a representative example. In particular, we focused on the automated determination of whether seeding activity induced by CWD prions exists in individual RT-QuIC reactions. We first analyzed a well-labeled, previously generated dataset using various AI models to assess the potential of AI in distinguishing true positive, false positive, and negative RT-QuIC reactions for CWD prions. We then evaluated and compared the performance of various AI models, including a widely used unsupervised algorithm (K-Means) and supervised models (SVM and MLP) (Figure 1b), on both summarized metrics and raw fluorescence data from RT-QuIC assays. Next, the best-performing model was applied to a more complex subset of the data not used during training, assessing whether AI can identify patterns in RT-QuIC data that are not readily discernible using current methods. The final testing consisted of testing the models on an independent validation dataset to evaluate generalizability of the models.

## Methods

### Data Generation and Processing

The dataset was sourced from Milstein, Gresch, et. al.^44^, a study that tested different disinfectants on CWD and consisted of the RT-QuIC analysis of swab samples after disinfection of various surfaces. The portion of the dataset used for this study comprised 8028 individual wells, representing 573 samples, each prepared with 2-3 dilutions and assayed in 4-8 technical replicates/reactions. Data from a total of 7331 sample reactions and 984 control reactions were used to evaluate the ML models. Among the 573 samples, 19 were withheld due to ambiguous labels, being classified as negative by statistical tests but positive by a "blind" human evaluation. These samples were further reviewed to exclude those with complex identities for additional characterization. The final dataset used for training and initial testing of the models included 7027 sample wells and 984 control wells. Each of these wells was characterized by taking a fluorescence reading 65 times at 45-minute intervals over a 48-hour run. Considering this sourcing, the ground-truth CWD positive/negative status of each sample was not directly known due to the introduction of disinfectant and the indirect nature of the swabbing method used to generate the RT-QuIC data (Figure 2). The "true" sample identities were instead determined by experienced researchers using statistical assessments, providing an excellent opportunity to compare the performance of AI solutions against existing methods for identifying CWD prions in RT-QuIC data.

**Fig. 2.**
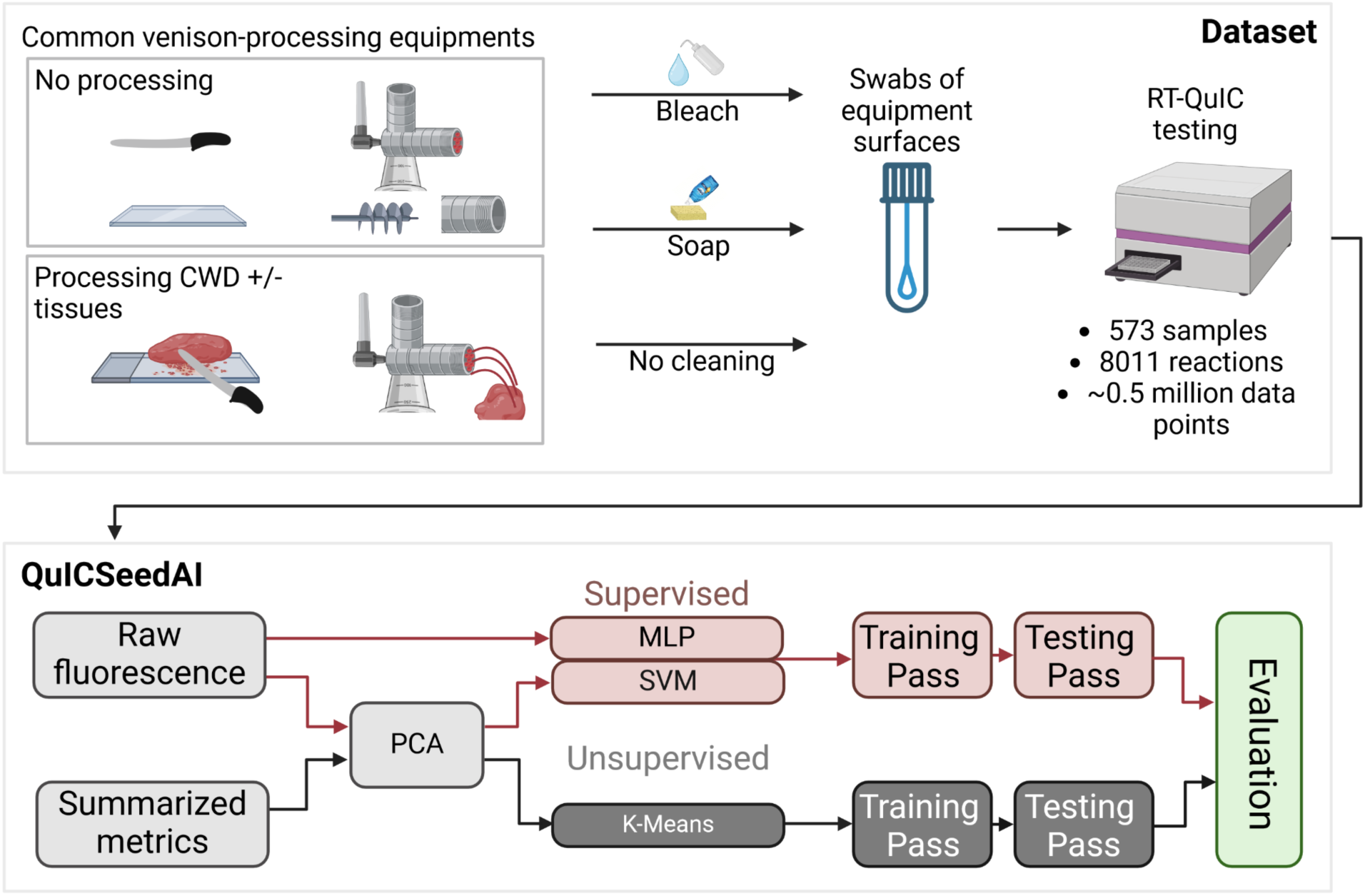
Visualization of Data Generation and Processing. Each reaction is obtained from the RT-QuIC analysis of CWD positive/negative samples obtained through the experimental design process outlined in the Milstein, Gresch, et. al. paper^44^. The fluorescence readings from this analysis are then processed into summarized metrics to be used as a comparative dataset. Both datasets, having been processed by PCA (except in the MLP) are split into a training and testing set. These datasets are used respectively to train and test the models, with the testing data providing the basis for evaluation.

Three labels were introduced for this study based on the results of each RT-QuIC reaction: negative, false positive, and true positive. Each of these labels is related to sample identity and the threshold defined in the original study^44^. A negative label represents a reaction involving a negative or a positive sample that never crossed the threshold (i.e. was identified as negative by analysis of the RT-QuIC results). A false positive label indicates a reaction with a CWD negative sample that crossed the threshold after being identified as a negative sample in the Milstein, Gresh, et. al. study^44^. A true positive label was identified as a reaction involving a CWD positive sample which crossed the positive threshold after being considered as positive in the original study^44^. In this dataset, there were 6781 negative reactions, 239 false positive reactions, and 991 positive reactions.

This formation of the dataset imposes a few limitations to the utility of the models evaluated in this study. Firstly, without knowing the identity or “ground truth” of the samples, these models can only be compared against the labels of annotators and, in the case of the supervised models, will be trained with any implicit biases held by the annotators. Additionally, the ambiguity in sample identity also limits the results; rather than classifying a reaction or sample as a whole, this method is limited to the classification of each replicate individually based on its annotation. While this individual replicate identity, or well-level identity, can inform sample identity, this limitation prevents the model from being an all-inclusive diagnostic tool. Despite this, the models will still learn meaningful patterns which could be invaluable to future sample-level identification and serve as a proof of concept for models that could potentially surpass current annotator/metrics-based classification methods.

In addition to this dataset, a smaller, external validation dataset was used to assert the validity of these methods by testing them on data generated completely independently from the training dataset. This data featured 122 negative reactions, 15 false positive reactions, and 31 positive reactions with 156 wells being sourced from samples and 12 being control reactions. For this dataset, the samples were derived directly from tissue samples (including ear, muscle, and blood), making the labels more reliable.^28,45,46^

Every method chosen for this study attempts to contribute to two goals: 1) to evaluate classification methods based on prevailing knowledge on the analysis of RT-QuIC data, and 2) to create a new method with the automation capabilities machine learning represents. These two goals were supported by two datasets—a dataset of precalculated metrics and a raw fluorescence dataset (Figure 1b). The first of these datasets, in line with the first goal of the study, consisted of four features extracted from the raw fluorescence values of each well. The feature set included the time to a precalculated threshold (TTT), the rate of amyloid formation (RAF), the maximum slope of the fluorescence curve (MS), and the maximum point ratio (MPR) between the smallest and largest fluorescence readings. These features were selected for their ability to characterize the curves RT-QuIC generates and for their relevance to the current method of using a precalculated threshold for classifying a test result as positive or negative. RAF and TTT are reciprocals of one another, with TTT measuring the “time required for fluorescence to reach twice the background fluorescence”^44^ and RAF being the rate at which this threshold is met. Both were included in the metrics dataset to give the feature selection algorithms used in this study the freedom to select whatever combination of these it deems best. Additionally, reactions which did not reach threshold were set to a TTT of 0 as this value is not physically realizable and works better for scaling than choosing some other large value which would need to depend on the runtime of the assay. MS was calculated with a sliding window with a width of 3.75 hours or 6 cycles. By finding the difference between the fluorescence reading at a current point and one taken 3.75 hours previously, the entire curve can be characterized into slopes. The maximum of these is selected. MPR was determined by dividing the maximum fluorescence reading by the background fluorescence reading.^44^ In contrast to this first, distilled dataset, the second dataset consisted of the raw fluorescence readouts, preserving all the information obtained from a particular run for the algorithms to extract. Once the two datasets were generated in the aforementioned formats, a few universal preprocessing steps were applied to ensure the data was ready for machine learning.

Firstly, RT-QuIC data in its raw form is not well suited to machine learning due to the wide variabilities in the absolute fluorescence. Different machines and concentrations, for example, can cause fluorescence curves to reach higher in some plates than in others, making it difficult to derive meaningful patterns in the larger dataset. The Scikit-Learn (sklearn) Standard Scaler^47^ was used to remedy this, centering the means of each well at 0 and scaling the dataset to unit variance. This was done separately to the training and testing partitions of the data. Secondly, as this study focused on the behavior of individual wells, it was crucial to remove any patterns in the data that could be derived from the ordering of the dataset. To account for this, the samples were shuffled into a random order using tools in the Numpy package.^48^

### Creation of Training/Testing Sets

For unsupervised models, it was advantageous to use the entire dataset for training, particularly on the feature-extracted dataset, as it demonstrates the success of classifying RT-QuIC data with only simple patterns. Without information from labels directing training, unsupervised models can learn an entire dataset with minimal risk of overfitting. The supervised models, however, required distinct training and testing sets to ensure the models could generalize beyond the training data. To accomplish this, 20% of the dataset was selected at random and withheld from training to be used in the evaluation of the models. The separation was done at the sample level to ensure different reactions from the same sample were not present in both the training and testing sets. The model predictions on this testing dataset are the source of the evaluation metrics and plots generated for each model. All of the models were configured to use the same data for training and testing to allow for direct comparison.

### Principal Component Analysis

Before training two of the model types, SVM and K-Means, a Principal Component Analysis (PCA) was applied to the two datasets. The PCA implementation used in this study was a part of the sklearn package^47^. PCA is a linear dimensionality reduction method that extracts relationships in a dataset and creates a new set of features based on these relationships. The algorithm works by identifying linear combinations of the different features in the dataset, matching each variable with every other variable and assigning a score based on how related they are to one another. This score is called the covariance. These scores are stored in a matrix which can be used to calculate eigenvectors which transform the original features into the set of linear combinations of features. These linear combinations are called Principal Components (PCs). The eigenvalues of these eigenvectors, then, represent the overall variance/feature importance of each individual feature. PCA can then select the features with the largest eigenvalues to represent the dataset.^49^ Typically the number of PCs to incorporate into the final dataset is identified by using a scree plot and identifying a point at which adding additional features has greatly reduced improvement in overall variance (also called finding the “elbow”). Using this method on the PCs obtained from the raw data, it was determined that 4 or 5 PCs were all that was necessary to represent the dataset. 4 PCs were selected from this set to be used in training some of the models as this aligned with the 4 features of the metrics dataset and captured nearly 90% of the variance in the dataset. The models which made use of PCA for their training dataset were the SVM and K-Means algorithms. Both of these algorithms train poorly on data with large numbers of features (such as the raw data with 65 timesteps used in this study), making PCA an effective way to give these algorithms an assist. Additionally, applying PCA to both datasets enhances the variance in the data, making the training of K-Means and SVM more efficient. In contrast, deep learning models, like the MLP, inherently handle high-dimensional data through their robust feature extraction capabilities^1^. Therefore, we did not apply PCA when training the MLP model, allowing it to learn directly from the raw data.

### Unsupervised Learning Approaches

#### K-Means Clustering

A K-Means model was trained on both the raw dataset as well as on the dataset of metrics, yielding a total of two models. KMeans is a powerful clustering utility and one of the most famous unsupervised learning algorithms. The algorithm works by identifying cluster centers through various methods and adjusting these locations/scales to optimally separate the dataset into distinct sets. The KMeans algorithm evaluated in this study was developed using the Scikit-Learn package in Python and used Lloyd’s KMeans Clustering Algorithm (LKCA)^50^. This algorithm uses an iterative optimization approach, and, like many machine learning algorithms, frequently finds solutions which are only locally optimal. Practically, this means that the solution KMeans will find to a clustering task is highly influenced by the initial conditions of the algorithm. In order to generate these initial conditions, the model was set to use a random initialization, selecting the features of two random samples as cluster centers which were iteratively adjusted by LKCA to find an optimal solution. LKCA was allowed to run up to 500 iterations or until the algorithm had converged on a solution - whichever required less time. Due to the high initialization sensitivity of KMeans, the algorithm was configured to select 150 random samples to use as initial cluster centers, selecting the best version. The goal was to greatly increase the odds that KMeans selected two samples, one which nearly epitomizes a negative sample and one which nearly epitomizes a positive sample. Then, with the optimization of LKCA, this increases the likelihood of finding a globally optimal solution. The raw and metric data were both passed through a PCA algorithm to highlight the variance in the data and reduce dimensionality.

K-Means, as an unsupervised model, can be configured to learn and cluster the entire dataset together, so the training and prediction steps both include all of the trainable data. The model was configured to find three clusters, with the idea that one cluster would represent positive RT-QuIC reactions, another would represent false positive reactions, and the final cluster would represent negative reactions. As most of the analysis used to evaluate the models was not compatible with multiclass output, the model output was converted to yield a 0 or a 1, corresponding to negative or positive. False positive classifications were treated as negative except when evaluating performance on false positives. Since the model was not given labels, it randomly selected whether the positive label was a 0 or a 1. The rest of the evaluation required a uniform system of outputs, so a standard was selected (1 represented positive and 0 represented negative). Any model that underperformed below what was predicted by random guessing had its labels flipped to match this standard..

### Supervised Learning Approaches

#### Support Vector Machines

While SVMs are elegant in their simplicity, they often suffer from poor performance on datasets with many features (as in our case with 65 timesteps). Considering this, the SVMs were considered prime candidates for the PCA, which was applied to both datasets via the Scikit-Learn package^47^ prior to training and testing. For the raw data, this created a much smaller feature space of just the relationships while enhancing the distinctions for the feature-extracted data. The SVM was constructed using the Scikit-Learn implementation^47^. The model itself uses a radial basis function (RBF)^51^ kernel. RBFs are a family of functions which are symmetric about a mean, such as a Gaussian distribution. These functions allow SVMs to classify data which is not separable by a polynomial function (such as concentric circles), making the SVM more versatile but also more complex^51^.

SVMs implemented with Scikit-Learn are inherently binary classifiers and do not support multiple classification boundaries. While it is possible to implement multiple classes by training multiple models, the implementation in this study did not use that approach in order to simplify the training and testing of the model. The PCA/SVM combo was run on the raw data and feature-extracted data.

#### Deep Learning: Multilayer Perceptron

Multilayer Perceptrons (MLPs) are one of the simpler types of Deep Neural Networks^1^ (DNNs) developed for classification tasks. MLPs consist of a set of 1-dimensional layers of neurons, featuring an input layer, an output layer, and at least one hidden layer in between. Each layer (with the exception of the input layer) learns to identify patterns in the previous layer which correspond to the training task^43^.

The MLP was selected for this task due to its natural compatibility with the dimensions of the problem. Since the study examined each replicate individually, the data was one-dimensional, consisting of just the fluorescence reading at each of the 65 timesteps. MLPs are able to compare each timestep with every other timestep, allowing them to extract a wide range of features from the dataset, even when these features are not linearly separable by class.

The MLP that produced the results in this paper used the Keras package in Python. Keras is a machine learning package which makes model development simpler and is part of the Tensorflow package^52^. The model consisted of 3 hidden layers. The input layer matched the input shape (65 nodes, one for each timestep). The next two hidden layers used 130 neurons each, which in testing allowed the model to learn the dataset better with more space for new features. The third hidden layer used 65 neurons as this limited overfitting in testing. Each of these layers used a Rectified Linear Unit (ReLU) activation function, a popular activation which eliminates negative inputs completely ^53^. The model, while originally tested as a binary classification setup, is configured with 3 output neurons, each corresponding to a class (negative, false positive, or positive). This forces the MLP to learn distinctions between positive and false positive classes in a more meaningful way. The model uses a softmax activation function in the final layer, converting the input to this layer into a score from 0 to 1 for each of the three classes. These scores add up to 1 and can be seen as the model’s “confidence” that a well belongs to a given class. With this information, the highest scoring class can be taken as the model’s prediction or can be processed more carefully to intuit new insights about the data.

Before the model is trained on the data, the inputs are first processed according to the steps outlined in the Data Generation and Processing section of the main manuscript. An important note, however, is that the MLP is the only model type explored in this study which does not require additional feature extraction. Neural networks are themselves feature extractors, so attempting to train the model on PCA extracted features or metrics would limit the potential features from which the model can extract information. This represents a massive advantage DNNs have over other ML methods as DNNs generally do not require time-consuming feature engineering to achieve good performance^1^.

Training an MLP involves passing training data through the network in chunks called batches. For this implementation, the number of reactions in a batch was 64. Once the network obtains predictions on this data, the prediction is compared to the true class using an error function (more commonly known as the loss function). This loss is then used to update the weights in between neurons of the network through a process called backpropagation ^1,54^. This process continues until all the training data has been passed through the network, completing one iteration or epoch of training. The MLP used in this study was set to train for a maximum of 100 epochs. While training, a portion of the training dataset is withheld (10% in this case) for validation of the model after each epoch. Should the model begin to overfit, a process in which the model loses the ability to generalize beyond the training data without learning more generally applicable patterns, the loss of the model on the validation data will begin to increase even as the training data loss decreases. The loss of the model on this validation set was monitored during training (Figure 3) and used to limit overfitting. At each epoch, the model was saved as a checkpoint if the validation loss at that epoch was less than the best previous validation loss. At the end of training, the last checkpoint was restored to the model iteration which performed best on the validation set.

**Fig. 3.**
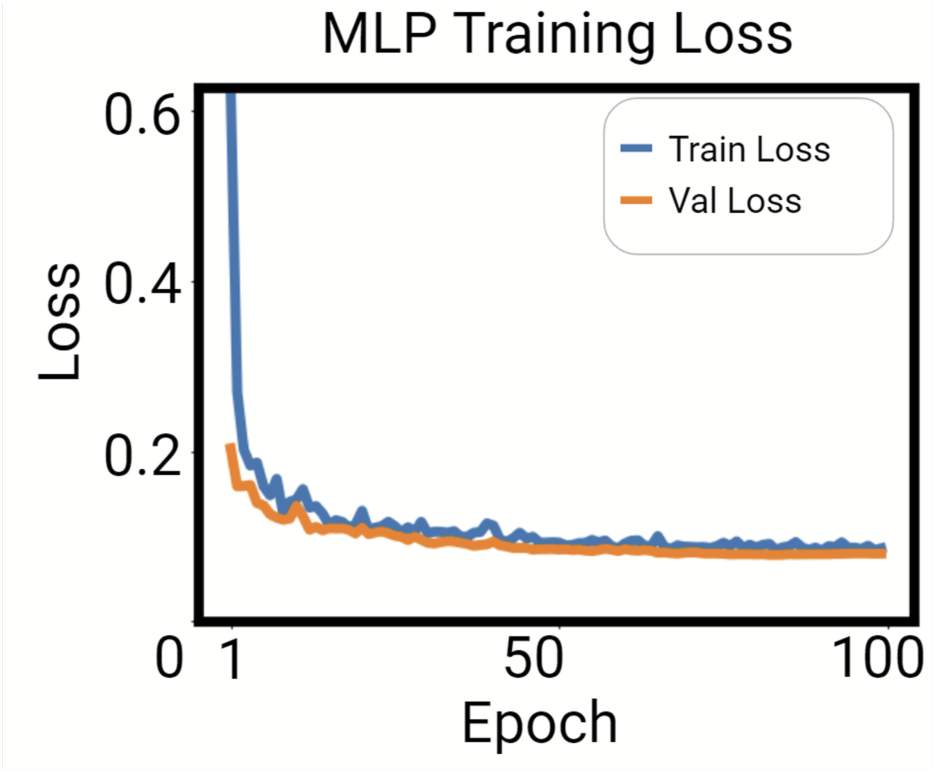
Multilayer Perceptron Loss. The value of the cross-entropy loss is measured after each epoch on the dataset used to train the model and a small portion of the larger training set which was withheld for validation. A lower loss indicates less error between predicted and true labels.

For the MLP used in this study, however, the model always improved on the validation set, so the checkpoint which was restored was the final state of the model after the 100th epoch.

Once the model was trained, the outright classification was a subpar representation of the data in this study. The false positive samples and the positive samples were not clearly distinguished by the model, meaning any well with a curve would score highly in both the false positive and positive class. This behavior was undesirable as many positives were mistaken for false positives and vice versa. In order to rectify this issue, a weighted average was applied to the output to get a single, unified score. The model output used the function shown in Eq. 1 to calculate the final score used for final classification. In the equation, Yneg is the score for the negative class, Yfp is the score for the false positive class, Ypos is the score for the positive class, and Yout is the overall binary score used in the results. This smoothed out the poorly defined class boundary into a spectrum from 0 to 2 (noting that the sum of the score outputs produced by the model are normalized), with any score less than 0.5 being labeled a negative prediction, any value between 0.5 and 1.5 being labeled a false positive prediction, and anything above 1.5 being labeled a positive prediction.

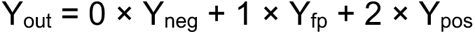

Equation 1: Weighted Average Applied to MLP Output

## Evaluation Criteria

Evaluation of the models primarily considered five metrics, accuracy, sensitivity/recall, specificity, precision, and F1-score. The accuracy metric corresponds to how many wells were classified the same as the human labels. As this study used 2 classes, the worst possible accuracy score should be 50% (when adjusted for class weight), corresponding to a random guess. A significantly lower accuracy would likely be the result of a procedural issue, so for each accuracy metric, it is imperative to consider the improvement over 50% as the true success rate of the model. While accuracy evaluates the dataset holistically, sensitivity (or recall) evaluates the performance of the model on identifying positive samples correctly. Specifically, sensitivity corresponds to the fraction of human-labeled positive wells that were classified correctly. Low sensitivity corresponds to a model that frequently classifies annotator-labeled positive samples as negative. Specificity, contrary to sensitivity, identifies the fraction of human-labeled negative wells that were classified correctly. Low specificity, then, correlates to a model that frequently classifies negative samples as positive. Another metric, precision, provides additional insight into how well a model is distinguishing between classes. Precision is defined as the fraction of model-labeled positive samples which were actually positive. This metric is of particular interest in this application as the number of negative reactions in the dataset is much greater than the number of positive reactions - an environment which can allow the previous metrics to be high despite a poorly performing model. In particular, precision measures a model’s false positive rate, with a low precision indicating a high rate of false positives relative to the number of true positives, making a positive prediction less meaningful. F1-score blends this idea of precision with sensitivity, creating a metric that evaluates the performance of a model on positive samples considering these factors together. While less useful for identifying patterns of mistakes in models, F1-score can be used to put sensitivity and precision in context together. F1-score is calculated as the harmonic mean of precision and sensitivity, meaning a particularly low precision or sensitivity will create a low F1-score.

## Results

### Extraction of early reaction features and validation of existing metrics by PCA

The PCA algorithm provided key insights into how the data can be distilled into only the most vital components. This information can then be used to generate insights into how the data is structured and, in the case that the identified variance relates to classification, can provide visuals of how clusters form.

This visual can be demonstrated in the form of a scatter plot (Figure 4) for both the raw data and the metrics data. In the case of the raw data, these two PCs account for nearly 80% of the variance in themselves, representing two of the four PCs used to train the raw K-Means and SVM models. In the case of the metrics data, these two PCs account for over 90% of the variance in the dataset and both represent linear combinations of the calculated metrics. In this case, it becomes clear that using all four of these metrics was unnecessary as TTT and RAF contain the same information. As such, only 3 PCs are required to represent the dataset in most cases. This would allow for a simplification of the models somewhat in the future. In addition to these insights, the two scatter plots also demonstrate the capability both datasets possess to meaningfully separate the data. Although there is overlap (particularly between the false positive and positive reactions), some distinction can be inferred between data points with different class identities.

**Fig. 4.**
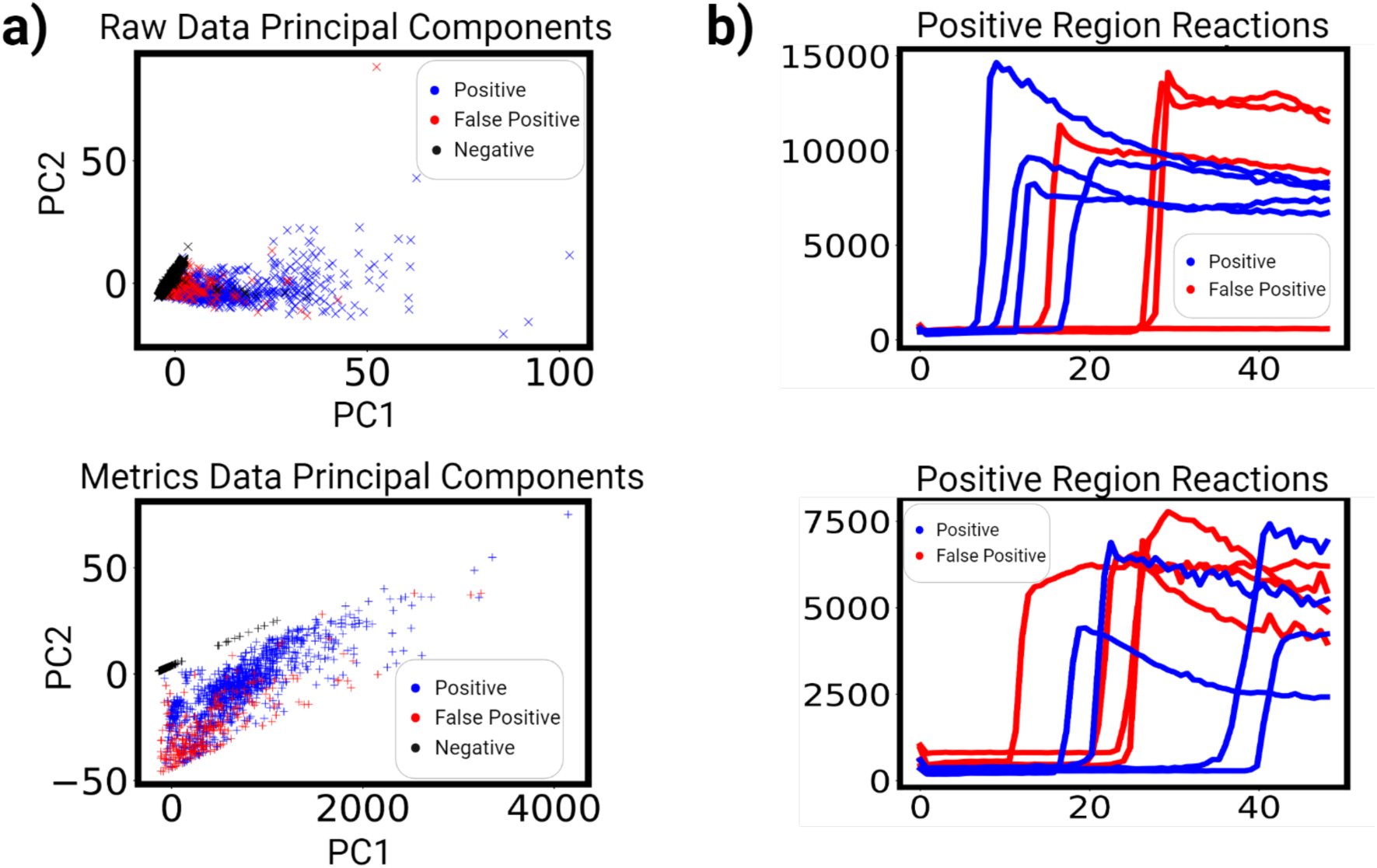
Demonstration of Useful Attributes Captured by PCA **a)** Class Separation Achieved by PCA. A PCA was run on the raw data and metrics data and generated principal components (PCs) corresponding to linear combinations of the input features. The figures demonstrate how these PCs create distinctions in the dataset which can be used with AI. **b)** Edge-case Examples. The PCA plots from a) were used to select the most positive-like false positives and compare with clearly defined positive samples. This demonstrates any likenesses which may make it difficult for PCA to create meaningful distinctions.

This is particularly interesting in the case of the metrics data, as this clear distinction indicates that the metrics (a significantly simpler dataset than its raw data counterpart) is sufficient to provide meaningful distinctions between classes. This validates the use of these existing metrics as a viable way of classifying RT-QuIC data. One oddity of this distribution, however, is the fact that all the negative samples appear to be closer to the positive samples than the false positives. This arises from the decision to set TTT for samples which never reached the threshold to a hard 0. PC1 and PC2 both depend on TTT and positive reactions will, in general, take less time to reach threshold than false positive reactions resulting from false seeding. Combining these PCs, then, causes this distinction to become apparent.

A few edge-case examples of positive and false positive reactions can also be plotted (Figure 4), providing a visualization of any reactions which may not be straightforward to classify. These edge-cases were identified from examination of the scatter plots and selection of data from each of these two classes which were furthest in the apparent positive region. Ultimately, these visualizations show the difficulty inherent to this dataset in the false positive reactions. Many of these reactions are difficult to distinguish from true positive reactions. Models with decent degrees of accuracy on these reactions, then, represent a powerful, standardized tool for handling edge cases without needing human annotation.

### K-Means metrics performed well for unsupervised RT-QuIC classification

Among classification algorithms, perhaps the most interesting and challenging to apply are unsupervised learning algorithms. Able to pick out patterns in unlabeled data, unsupervised algorithms provide a critical foundation in understanding the dynamics of a dataset, allowing us to determine if the variances PCA and metrics-based feature extraction identifies are meaningful to classifying RT-QuIC reactions. The patterns the algorithms select are not directed as in supervised learning, so good performance of these models indicates a dataset in which the predominant patterns are the ones pertaining to the classification of RT-QuIC reactions as positive or negative.

#### K-Means

The model performance metrics for the PCA feature-extracted dataset and for the manually feature-extracted dataset were calculated (Table 1). For the raw data, the results represent an accuracy of 93%. For the manually feature-extracted data, the results represent an accuracy of 97%.

**Table 1:**
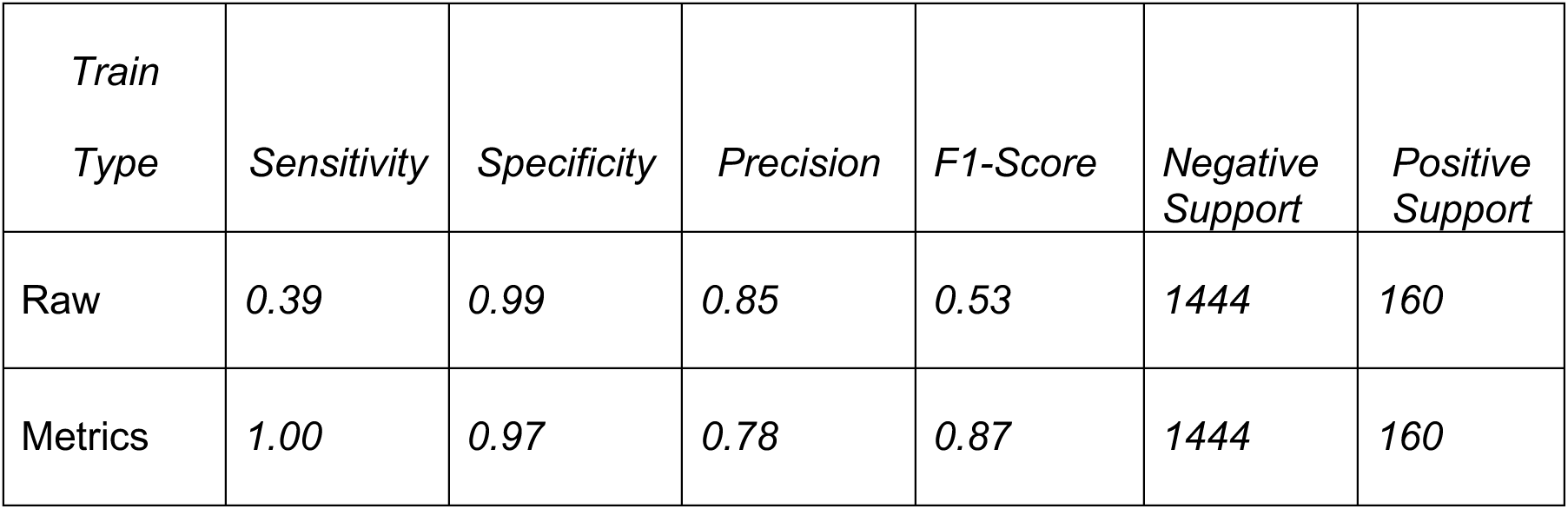
K-Means Performance Metrics (Higher is Better)

The K-Means model trained on the raw data learned to classify many positive samples as negative, implying that the learned cluster centers are skewed in an undesired way. The K-Means model trained on metrics, however, achieved high accuracy and excellent sensitivity. This suggests that the metrics provided a more useful basis for extracting salient features for the K-Means model to use.

### Supervised learning achieved high accuracy for RT-QuIC classification

Supervised algorithms, having access to labels during training, provide an important point of comparison against unsupervised models for learning patterns in datasets. These algorithms excel at finding patterns that support class identities specified by the operator, making them more adept at classifying more complex datasets.

#### Support Vector Machine

The SVM results represent 93% accuracy on the raw data as well as 98% accuracy on the metrics data (Table 2).

**Table 2:** SVM Performance Metrics (Higher is Better)

| *Train Type* | *Sensitivity* | *Specificity* | *Precision* | *F1-Score* | *Negative Support* | *Positive Support* |
| --- | --- | --- | --- | --- | --- | --- |
| Raw | 0.94 | 0.93 | 0.59 | 0.72 | 1444 | 160 |
| Metrics | 0.93 | 0.98 | 0.84 | 0.88 | 1444 | 160 |

The SVM performed very well on the testing data and with minimal training time, with the metrics trained model outperforming the raw data/PCA approach in most metrics. The high accuracy of this model compounded with the highest precision of any of the models evaluated in this study makes this a compelling choice. The sensitivity is also the lowest out of all the models, however, incurring a cost for the high precision.

#### Multilayer Perceptron

The metrics produced by the MLP were calculated and represent an accuracy of 97% (Table 3).

**Table 3:** MLP on Raw Data Metrics (Higher is Better)

| *Train Type* | *Sensitivity* | *Specificity* | *Precision* | *F1-Score* | *Negative Support* | *Positive Support* |
| --- | --- | --- | --- | --- | --- | --- |
| Raw | 0.98 | 0.97 | 0.80 | 0.88 | 1444 | 160 |

**Tab. 4.** Replicate Reactions in Agreement with Human Annotation.

|  | SVM Metric | MLP | K-Means Metrics |
| --- | --- | --- | --- |
| Reactions Agreed with Human annotation | 304/304 | 292/304 | 304/304 |

Requiring less preprocessing than the SVMs and K-Means, the MLP benefited from a simpler implementation (feature-engineering-free) while still achieving high accuracy and specificity. This positions the MLP as an effective and adaptable model that is not significantly affected by specific implementation. The MLP appeared to be an excellent all-around candidate for classification, maintaining comparatively high precision without a major sacrifice to sensitivity. Leveraging its high performance compounded with its versatile implementation, the MLP represents the capability of deep learning to be broadly applied without the need for expert-dependent feature engineering. This makes the MLP and other deep learning techniques prime candidates for classification of SAA data generated using other neurodegenerative diseases without the need for application-specific feature extraction.

*Summary.* The supervised learning methods used in this study represent a useful alternative to the unsupervised learning methods presented previously. Both the SVM trained on metrics and the MLP achieved relatively higher precision than the unsupervised models, making them more useful in applications where negative wells greatly outnumber positive wells. Particularly in the case of the MLP, deep learning was able to identify useful relationships in the dataset with a relatively simple implementation and achieved excellent all-around results. This highlights the capability of DNNs to extract salient features from data without the need for expert feature engineering, making these networks powerful choices for datasets which have not been well characterized.

### Comparative analysis of unsupervised and supervised learning for RT-QuIC classification revealed key performance differences

#### Unsupervised Overview

The confusion matrices (Figure 5a) and ROC curves (Figure 5b) highlight the story told by the tables previously in greater clarity. The K-Means trained on the metrics data is the best performer of the two, never missing a positive sample in the entire testing dataset. Additionally, the 3.6% of negative samples it classified as positive are made up entirely of false positives.

**Fig. 5.**
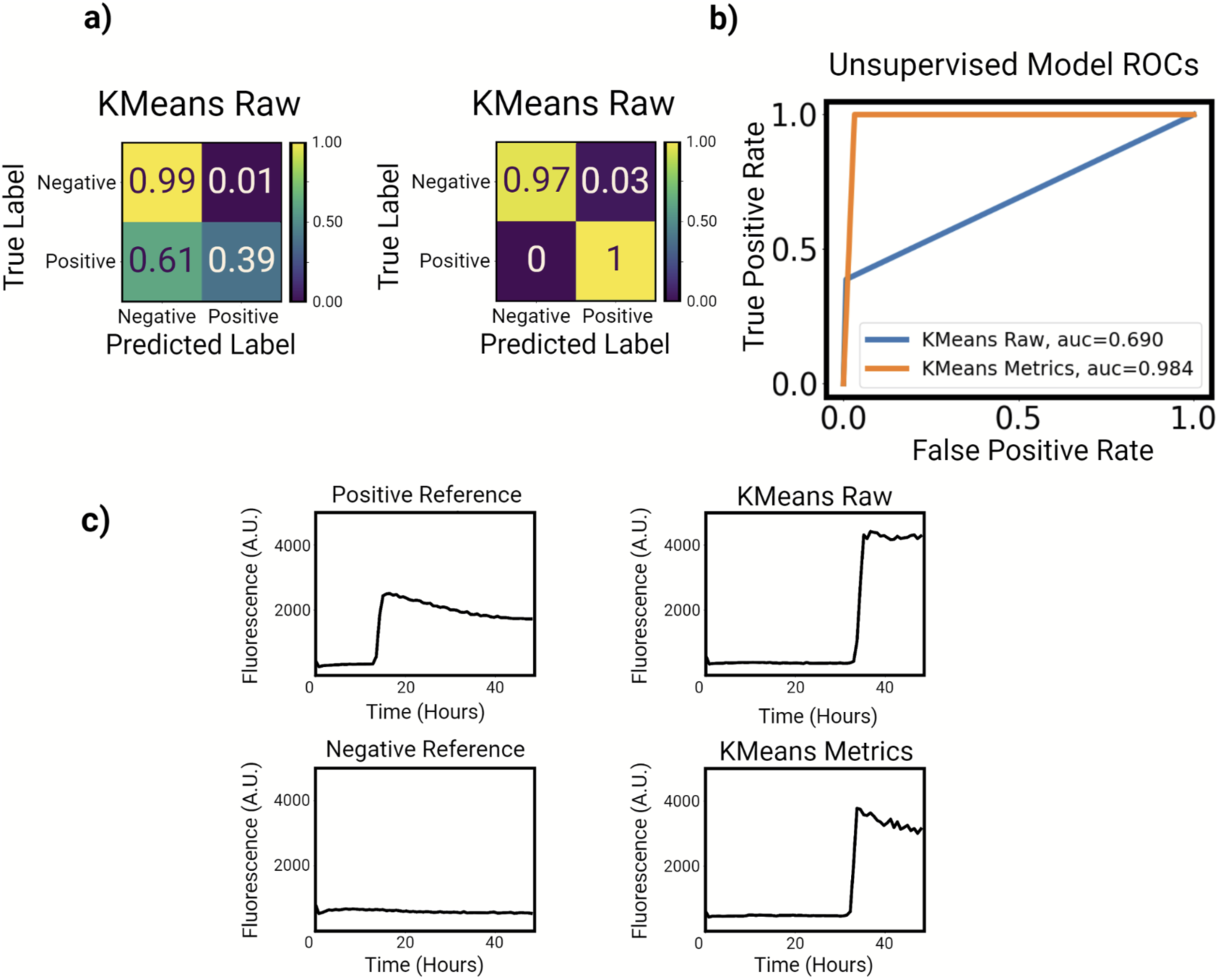
Comparison of K-Means model results. **a)** Model confusion matrices represent the ratio of how many samples from each true class were classified in each predicted class. **b)** The Receiver Operator Characteristic (ROC) illustrates the sensitivity-specificity tradeoff of each model, with an ideal model having an area under the curve (AUC) of 1. **c)** The examples include a single false positive a given model misclassified along with correctly classified reference samples, highlighting how distinguishable a misclassified false positive was.

This model never successfully classified a false positive and the false positives are the only reactions the model misclassified. The example curves (Figure 5c) also support the finding in the PCA analysis that these false positive curves are not always trivial to classify.

#### Supervised Overview

In the case of the supervised learning methods, while there is a clear worst model, there is no clear best model between the SVM trained on metrics and the MLP. Both of these models achieved different goals with different tradeoffs in approach. In the case of the MLP, while the results were more balanced, the slightly worse performance on correctly identifying negative reactions represents a much larger portion of the dataset as negative reactions are more common in general. This higher prioritization of correct classification of positive samples, however, favors the MLP in the ROC evaluation (Figure 7b). As was the case with the unsupervised models, the example plots demonstrate the challenge of classifying some of the false positive samples (Figure 6c, Figure 7c).

**Fig. 6.**
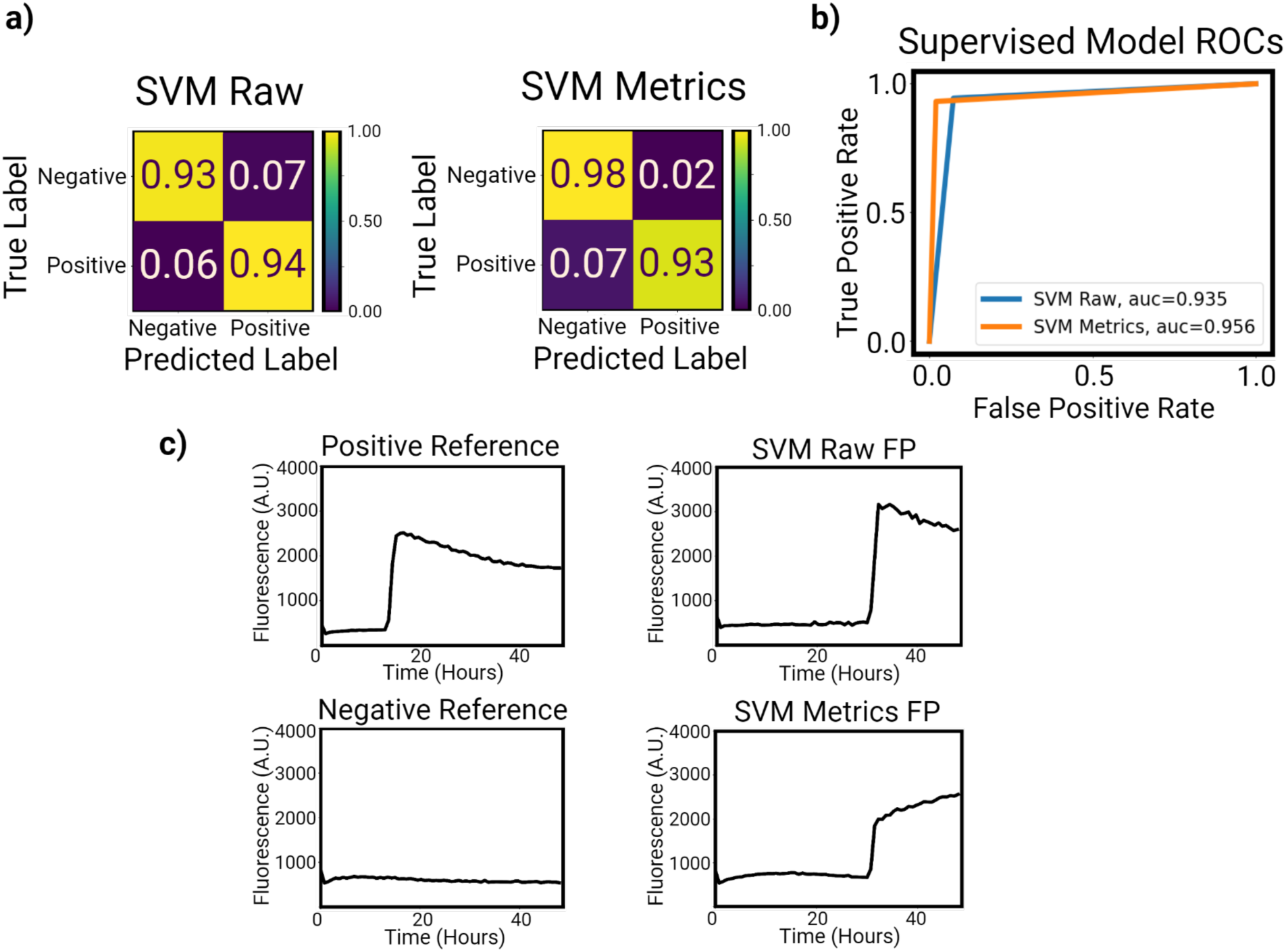
Comparison of SVM model results. a) Model confusion matrices represent the ratio of how many samples from each true class were classified in each predicted class. b) The Receiver Operator Characteristic (ROC) illustrates the sensitivity-specificity tradeoff of each model, with an ideal model having an area under the curve (AUC) of 1. c) The examples include a single false positive a given model misclassified along with correctly classified reference samples, highlighting how distinguishable a misclassified false positive was.

**Fig. 7.**
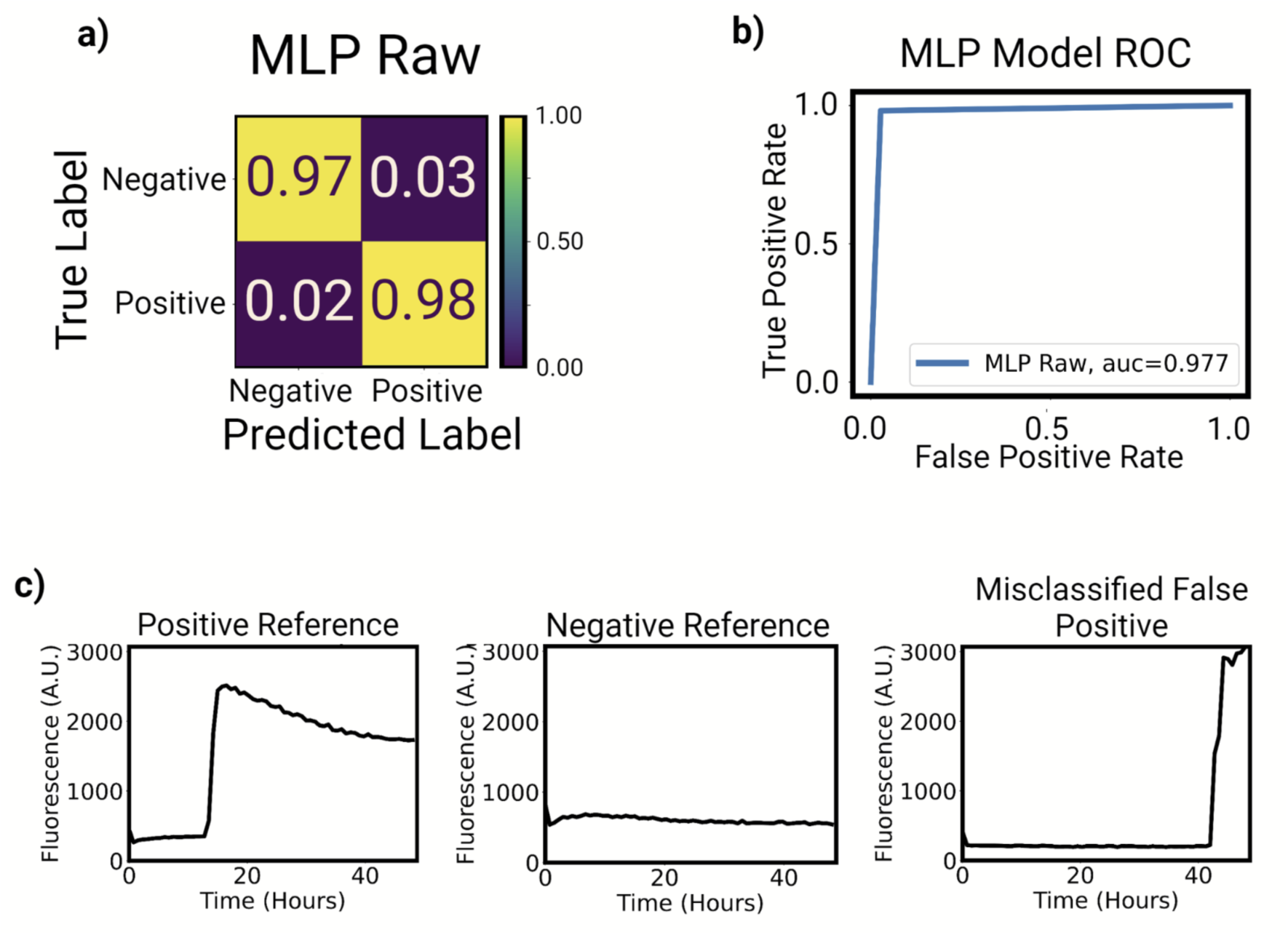
MLP model results **a)** Model confusion matrix represents the ratio of how many samples from each true class were classified in each predicted class. **b)** The Receiver Operator Characteristic (ROC) illustrates the sensitivity-specificity tradeoff of the model, with an ideal model having an area under the curve (AUC) of 1. **c)** The examples include a single false positive a given model misclassified along with correctly classified reference samples, highlighting how distinguishable a misclassified false positive was.

#### Performance on Dataset with Label Ambiguity

In addition to the 554 samples which made up the training and testing dataset for the models, 19 samples with ambiguous labels were tested separately. This subset of the dataset consists of swab samples where human annotators, using a plot of the replicates overlayed for each swab, classified the swab sample differently than what was suggested by statistics. In particular, each of these samples was identified as positive by a human annotator, but determined to be negative by statistical analysis in the original study^44^. As a point of comparison, the two best-performing models were selected (the K-Means model trained on metrics and the MLP) and tested on these samples.

Each sample dilution was annotated by the K-Means model trained on metrics, the MLP, and a human annotator (ML). Examining these reactions, it becomes clear that machine learning represents a far superior method of classification to other statistical methods of evaluation. The excellent agreement between the models and human annotation represents a major improvement and demonstrates the capability of AI to limit the need for human annotation, providing a standardized, automated approach with accuracy approaching that of humans.

### Models performance on external validation data shows generalizability

In addition to the testing and evaluation performed on a subset of the original dataset, each model was applied to an external validation dataset^28,45,46^ to highlight generalizability. The results of this evaluation (Figure 8) demonstrate the capabilities of the different models to generalize to this new dataset. With the exception of the KMeans model trained on raw data, each model performed generally well on this dataset. One important distinction arises when comparing the supervised and unsupervised models, however. Each of the unsupervised models generalized poorly, losing significant performance on this validation dataset. KMeans Metrics, in particular, lost its 100% sensitivity for a sensitivity closer to 87%. The supervised models all performed similarly, however, with the raw data SVM achieving the highest specificity while the MLP and SVM trained on metrics both achieving similar performance with near perfect sensitivity. With the smaller size of this dataset, it is difficult to pick a clearly best generalized model. The supervised models, however, were far superior at generalizing than the unsupervised models.

**Fig. 8.**
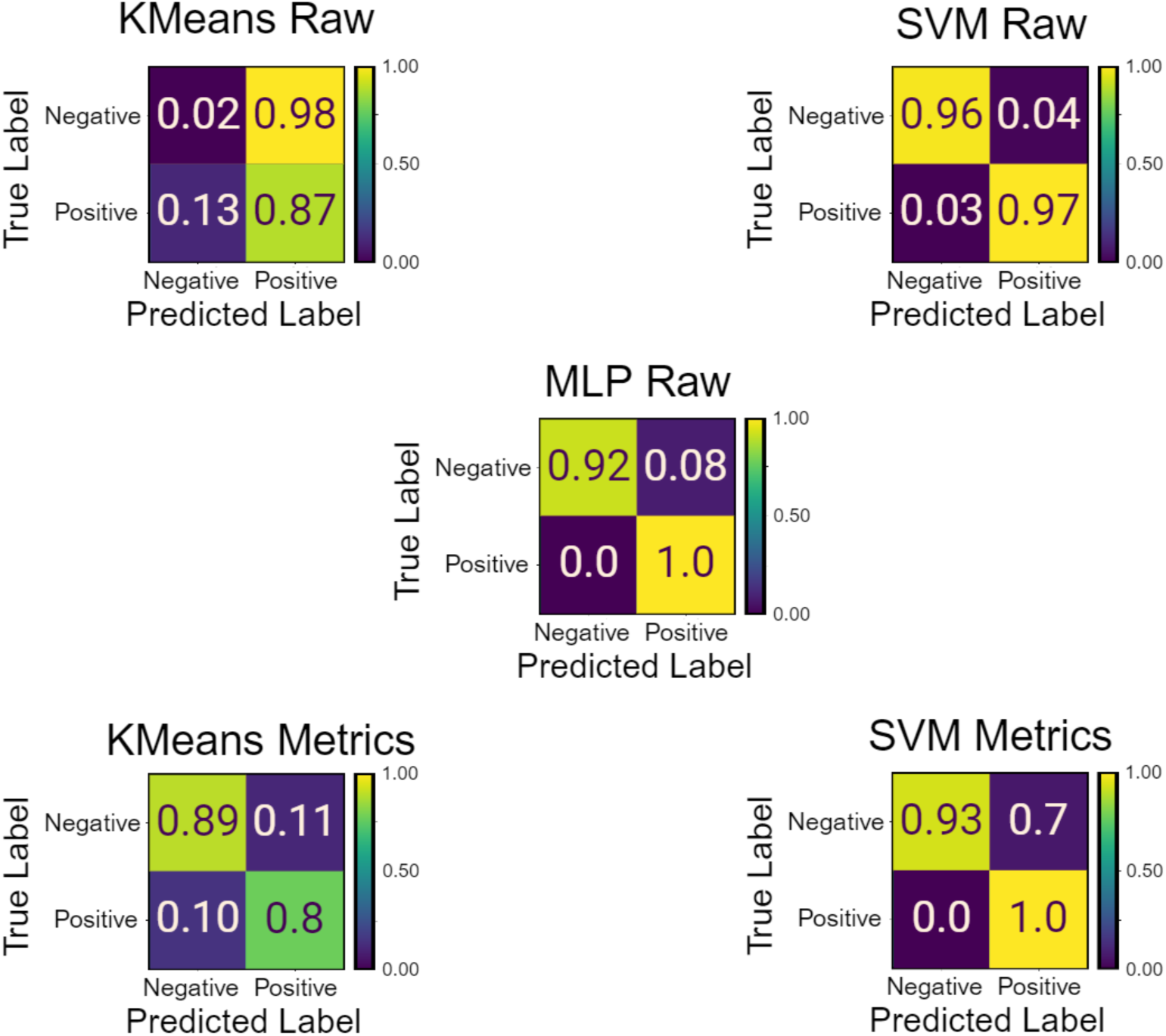
Model Results on External Validation Data. Model confusion matrices represent the ratio of how many samples from each true class were classified in each predicted class.The data used to generate these confusion matrices was generated independently from the rest of the dataset.

## Discussion

This proof-of-concept study demonstrated the promising potential of AI in enhancing the interpretation and automation of RT-QuIC data, using Chronic Wasting Disease (CWD) as a robust model for protein misfolding disorders. The results addressed two primary objectives: first, our comparative analysis of unsupervised (K-Means) and supervised (SVMs, MLPs) AI models unveiled their unique strengths in processing both summarized metrics and raw fluorescence data; second, we established that AI models can effectively identify seeding activity in RT-QuIC reactions.

Our findings demonstrate that AI models can effectively automate the detection of misfolded protein seeding activity in RT-QuIC assays. Notably, the deep learning-based MLP approach performed similarly to or better than other models by leveraging its inherent ability to extract relevant features automatically, directly learning from raw data without requiring dimensionality reduction techniques like PCA. This eliminates the need for expert-dependent and time-consuming feature engineering, streamlining the analysis process. The MLP model possesses adaptability to diverse datasets and assay conditions. By reducing reliance on data preprocessing, the MLP approach offers a more efficient and scalable solution for standardizing SAA data interpretation across various neurodegenerative diseases.

Collectively, our study lays the groundwork for AI-driven enhancements in RT-QuIC data analysis and the development of AI-assisted diagnostic tools for a spectrum of neurodegenerative disorders characterized by protein misfolding.

### Evaluation of Metrics

In examining the capabilities of the various models and comparing performance between the two datasets, a pattern of support emerges for the metrics selected to describe the curves. Firstly, it is worth acknowledging the apparent similarities between the PCA-selected features and the chosen metrics. The analysis of the PCA (Figure 4) demonstrates evidence that machine learning and human intuition for curve description are both capable techniques for clustering data. As is the case with nearly all machine learning applications, the exact patterns identified by the algorithm do not perfectly line up with human-identified patterns and concepts.

### Automation of Well Identification

The other primary goal of this study was to provide a method of reliably automating curve identification. Identifying a curve as positive or negative by the human eye can be time-consuming and inconsistent among labs and technicians. As such, AI represents a powerful solution, being highly adaptable and capable of identifying key patterns without significant technician time. While a few models stood apart as particularly effective at classifying the data in this study, nearly all of them demonstrate the viability of this approach, obtaining high accuracy similar to that of a technician. In addition to this high accuracy, the models are all highly reproducible and consistent. Once the dataset has been generated and the code written, these models each can classify tens of thousands of samples in seconds.

### Future Directions

Artificial intelligence (AI), particularly machine learning (ML), has shown remarkable effectiveness in identifying patterns that hold predictive value. Our study utilized the dataset of over 8,000 RT-QuIC reactions, the largest of its kind to contain both human annotations and metrics commonly used on RT-QuIC data, providing unprecedented depth for analysis^44^. With research efforts involving more comprehensive datasets, various misfolded proteins, and the development of AI models specifically tailored for SAA data, these models could potentially match or surpass human experts in distinguishing positive from negative samples, overcoming current challenges such as strain differences, cross-seeding, and concentration effects that could hinder accurate diagnosis in clinical settings. Future integration of AI with SAAs holds the potential to enhance diagnostic accuracy, streamline workflows, and ensure consistent interpretation across various assays and laboratory conditions, mitigating discrepancies in interpretation among personnel.

The implications of our findings extend beyond CWD and RT-QuIC to other SAAs, such as PMCA^21,23^, HANABI^55^, Nano-QuIC^32,56^, Cap-QuIC^57^, MN-QuIC^33^, which face similar challenges in data interpretation. AI-driven approaches offer the potential for standardization and automation across a spectrum of neurodegenerative diseases that have already shown seeding activity detectable by SAAs, including Alzheimer’s^10,21,58,59^, Parkinson’s^23,60^, as well as ALS and frontotemporal dementia^25,61^.

Looking forward, we propose the continued development of AI-QuIC to integrate diverse data types such as image and spectral data, and accommodate a wider range of assay parameters, enhancing its applicability to numerous neurodegenerative diseases. In addition, forecasting techniques and in-depth time-series analysis could lay the foundation for reduction of runtimes of the RT-QuIC assay, increasing speed and accuracy in an all-inclusive diagnostic framework. While the utilization of AI-QuIC on RT-QuIC data in this study is in its early stages, it represents a significant advancement toward the early detection and diagnosis of neurodegenerative diseases. By harnessing the pattern-recognition capabilities of AI - particularly the streamlined, feature-engineering-free approach of the deep learning MLP model - AI-QuIC effectively captures the dynamics of protein misfolding during SAA assays, enabling automated and standardized data interpretation.

Just as AI has revolutionized protein folding modeling, the integration of AI into SAA analysis shows the potential to enhance diagnostic workflows and contribute to drug discovery pipelines. Recent breakthroughs such as the cryo-electron microscopy determination of the atomic structure of misfolded prions^62^ have unveiled the structures of misfolded protein aggregates, providing new insights into their pathogenic mechanisms. These structural data also offer training data for AI to better predict not only normal protein folding but also misfolding processes. As AI continues to advance in modeling both protein folding and misfolding, it may facilitate the identification of new drug targets for protein misfolding diseases. By combining AI-enhanced diagnostics such as AI-QuIC with AI-based modeling of protein misfolding, we can accelerate our understanding of neurodegenerative disorders and ability to diagnose them at an early stage.

## Data availability statement

Data used in this manuscript may be obtained from the authors upon reasonable request.

## Acknowledgments

Portion of this research was supported by the Minnesota State Legislature through the Minnesota Legislative-Citizen Commission on Minnesota Resources (LCCMR) and Minnesota Agricultural Experiment Station Rapid Agricultural Response Fund to K.D.H., M.L., P.R.C., P.A.L. and S.-H.O. M.L., P.A.L., and S.-H.O. acknowledge support from the Minnesota Partnership for Biotechnology and Medical Genomics. S.-H.O. further acknowledges support from the Sanford P. Bordeau Chair and the International Institute for Biosensing (IIB) at the University of Minnesota. The figures in this manuscript were created with the assistance of Biorender.

**Author contributions**

S.-H.O. conceived the study. M.L. and K.D.H. designed the study. K.D.H., M.L., S.-H.O. wrote the manuscript based on the input from other co-authors. M.L. curated data for the study. K.D.H. conducted the analysis utilized in the study. K.D.H., M.L., and P.R.C. visualized the data. K.D.H., M.L., P.R.C., P.A.L., and S.-H.O. discussed the results. All authors revised the manuscript. All authors approved the manuscript.

