## Supporting Information for "AI-QuIC: Machine Learning for Automated Detection of Misfolded Proteins in Seed Amplification Assays"

| No Dillution | | | | 10-fold Dilution | | | | |
|---|---|---|---|---|---|---|---|---|
| Sample: | MLP: | KMeans M: | Human: | | MLP: | KMeans M: | Human: | Number of Replicates |
| 49G | 4 | 4 | 4 | | 0 | 0 | 0 | 8 |
| 14G | 1 | 1 | 1 | | 1 | 1 | 1 | 8 |
| 92G | 0 | 0 | 0 | | 5 | 5 | 5 | 8 |
| 98G | 0 | 0 | 0 | | 4 | 4 | 4 | 8 |

| | | | | | | | | |
|---|---|---|---|---|---|---|---|---|
| 104G | 1 | 1 | 1 | | 6 | 6 | 6 | 8 |
| 121G | 4 | 4 | 4 | | 3 | 3 | 3 | 8 |
| 152G | 0 | 1 | 1 | | 4 | 4 | 4 | 8 |
| 157G | 3 | 3 | 3 | | 4 | 4 | 4 | 8 |
| 158G | 2 | 2 | 2 | | 4 | 4 | 4 | 8 |
| 205G | 1 | 5 | 5 | | 1 | 2 | 2 | 8 |
| 206G | 1 | 5 | 5 | | 0 | 0 | 0 | 8 |
| 212G | 0 | 0 | 0 | | 4 | 4 | 4 | 8 |
| 217G | 3 | 3 | 3 | | 2 | 2 | 2 | 8 |
| 267G | 0 | 0 | 0 | | 5 | 5 | 5 | 8 |
| 379G | 3 | 3 | 3 | | 0 | 0 | 0 | 8 |
| 319G | 4 | 4 | 4 | | 3 | 3 | 3 | 8 |
| 368G | 1 | 1 | 1 | | 4 | 4 | 4 | 8 |
| 321G | 0 | 0 | 0 | | 4 | 4 | 4 | 8 |
| 325G | 0 | 0 | 0 | | 3 | 4 | 4 | 8 |

Tab. 1 A Comparison of MLP, K-Means Metrics, and Human Annotations of Ambiguously Labeled Samples

| No Dilution | | | | 10-fold Dilution | | | | |
|---|---|---|---|---|---|---|---|---|
| Sample: | SVM Raw: | SVM M: | Human: | | SVM Raw: | SVM M: | Human: | Number of Replicates |
| 49G | 3 | 4 | 4 | | 0 | 0 | 0 | 8 |
| 14G | 0 | 1 | 1 | | 0 | 1 | 1 | 8 |
| 92G | 0 | 0 | 0 | | 4 | 5 | 5 | 8 |
| 98G | 0 | 0 | 0 | | 2 | 4 | 4 | 8 |
| 104G | 0 | 1 | 1 | | 2 | 6 | 6 | 8 |
| 121G | 4 | 4 | 4 | | 2 | 3 | 3 | 8 |
| 152G | 0 | 1 | 1 | | 4 | 4 | 4 | 8 |
| 157G | 1 | 3 | 3 | | 3 | 4 | 4 | 8 |
| 158G | 3 | 2 | 2 | | 0 | 4 | 4 | 8 |
| 205G | 1 | 5 | 5 | | 1 | 2 | 2 | 8 |
| 206G | 1 | 5 | 5 | | 0 | 0 | 0 | 8 |
| 212G | 0 | 0 | 0 | | 4 | 4 | 4 | 8 |
| 217G | 3 | 3 | 3 | | 2 | 2 | 2 | 8 |
| 267G | 0 | 0 | 0 | | 1 | 5 | 5 | 8 |

| | | | | | | | | |
|---|---|---|---|---|---|---|---|---|
| 379G | 3 | 3 | 3 | | 0 | 0 | 0 | 8 |
| 319G | 3 | 4 | 4 | | 2 | 3 | 3 | 8* |
| 368G | 0 | 1 | 1 | | 2 | 4 | 4 | 8 |
| 321G | 0 | 0 | 0 | | 3 | 4 | 4 | 8 |
| 325G | 0 | 0 | 0 | | 3 | 4 | 4 | 8 |

Tab. 2 A Comparison of SVM Raw, SVM Metrics Metrics, and Human Annotations of Ambiguously Labeled Samples
